# Activation of TPC2 amplifies lysosome-mitochondria calcium transfer to regulate energetic stress responses

**DOI:** 10.64898/2026.03.03.709381

**Authors:** Sadia Ahmed, Namratha Javvaji, Katherine L. Hammond, Kevin M. Casin, John W. Elrod, Paul M. Holloway, Yvonne Couch, Jillian N. Simon

## Abstract

Mitochondrial Ca^2+^ uptake governs metabolism and cell fate, yet how signals from other organelles shape this remains incompletely defined. Although lysosomes are relatively small Ca^2+^ stores, their strategic positioning at organelle contact sites suggests they may amplify Ca^2+^ transfer within nanodomains. Here, we show that activation of the lysosomal Two-pore channel 2 (TPC2) initiates rapid mitochondrial Ca^2+^ uptake through an endoplasmic reticulum-dependent relay requiring IP₃ receptors and the mitochondrial calcium uniporter channel. The extent of mitochondrial Ca^2+^ accumulation scales with TPC2 activity without affecting global Ca^2+^ responses, identifying TPC2 as a specific amplifier of lysosome-mitochondria Ca^2+^ exchange. Moderate TPC2 activation transiently enhances oxidative phosphorylation, whereas sustained enhancement increases susceptibility to Ca^2+^-induced mitochondrial permeability transition. In stroke models, hyperactivation of TPC2 exacerbates injury, while acute pharmacological inhibition at reperfusion confers neuroprotection, including in human iPSC-derived neurons. Thus, lysosomal Ca^2+^ release acts as an upstream regulator of mitochondrial energetic resilience under stress.

## Introduction

Mitochondrial Ca^2+^ handling plays a critical role in maintaining cellular homeostasis. Ca^2+^ uptake into mitochondria supports oxidative phosphorylation (OxPhos), buffers cytosolic Ca^2+^ transients, and regulates pathways central to cell survival and death.^1,2^ Dysregulation of this process is implicated in a broad range of pathologies, including cardiovascular disease^3,4^, skeletal muscle disorders^5^, and cancer^6^, underscoring the importance of tightly controlled mitochondrial Ca^2+^ flux. Furthermore, mitochondrial Ca^2+^ dynamics are crucial for maintaining neuronal health, and dysregulated Ca^2+^ homeostasis in the brain has been implicated in a variety of neurodegenerative conditions^7–9^ and in ischemia-reperfusion (I/R) injury^10,11^ where altered mitochondrial function causally contributes to cell death and tissue damage.

Recent studies have highlighted the importance of inter-organelle communication in regulating mitochondrial Ca^2+^ levels. Mitochondria do not function in isolation; rather, they are an important signaling hub that dynamically interact with various organelles, such as the endoplasmic reticulum (ER), lysosomes, and the plasma membrane, at specialized contact sites^1,12^. While the ER is the principal Ca^2+^ store^1^, emerging studies indicate that lysosomes also contribute to mitochondrial Ca^2+^ regulation.^13,14^ Prior work has characterized mechanisms by which lysosome-derived Ca^2+^, particularly via Transient receptor potential mucolipin 1 (TRPML1), influences mitochondrial dynamics and morphology.^13^ However, key questions remain regarding how lysosomal Ca^2+^ signaling, through alternative channels such as Two-pore channel 2 (TPC2), integrates with canonical ER–mitochondria pathways, and how the magnitude and timing of lysosome-mediated Ca^2+^ flux modulate mitochondrial function, bioenergetics, and cellular stress responses. Addressing these gaps is essential to understanding the lysosome’s role in multi-organelle Ca^2+^ crosstalk.

To address these questions, it is important to consider the unique properties of lysosomes that enable them to modulate mitochondrial function. Although lysosomes occupy only 1–2% of total cell volume, they contain high concentrations of free luminal Ca^2+^ (∼300–600 μM)^15^. Within spatially confined nanodomains, lysosomes can locally elevate Ca^2+^ levels, exerting substantial signaling effects.^16–19^ Through TPC2, lysosomes release Ca^2+^ into the cytosol in response to a variety of intercellular cues,^16,20^ influencing nearby organelles such as mitochondria.^18,21^ Yet, while prior studies have established lysosomal Ca^2+^ release as a regulator of mitochondrial dynamics, the full extent of its role in mitochondrial Ca^2+^ uptake, bioenergetics, and cellular stress resilience remains to be explored.

Here, we show that lysosomal TPC2 mediates rapid Ca^2+^ transfer to mitochondria through ER and mitochondrial Ca^2+^ uptake pathways. We further demonstrate that TPC2 activity is tunable, with moderate activation enhancing mitochondrial function, while hyperactivation leads to Ca^2+^ overload and increased susceptibility to stress. Using both in vitro and in vivo stroke models, we find that excessive lysosome–mitochondria Ca^2+^ exchange directly contributes to worsened neuronal injury, whereas pharmacological TPC2 inhibition affords neuroprotection. These findings establish TPC2 as a previously unrecognized regulator of mitochondrial Ca^2+^ signaling, highlighting a multi-organelle axis that could be targeted to modulate cellular resilience.

## Results

### TPC2 activation induces rapid mitochondrial Ca^2+^ uptake

To investigate the relative contributions of lysosomal Ca^2+^ channels to mitochondrial Ca^2+^ dynamics in excitable cells, we compared TRPML1 and TPC2 agonists using live-cell imaging in neuroblastoma N2A cells expressing the mitochondrial matrix-targeted genetic calcium sensor, mito-R-GECO. While previous studies demonstrated that TRPML1 activation can enhance local mitochondrial Ca^2+^ at lysosome–mitochondria contact sites,^13^ the impact on global mitochondrial Ca^2+^ uptake and function remains unexplored. Representative mito-R-GECO images are shown in **Fig. 1A**, with corresponding traces and quantifications in **Fig. 1B–G**. Consistent with minimal lysosome-mitochondria coupling through TRPML1 in this context, stimulation with the TRPML1 agonist, ML-SA1, failed to elicit a detectable increase in mitochondrial matrix Ca^2+^ over a 5-minute recording (F/F₀ = 1.08 ± 0.19 vs. 0.93 ± 0.10 in vehicle; **Fig. 1C**). In contrast, activation of TPC2 using the NAADP analog, TPC2-A1-N, triggered a rapid and pronounced rise in mitochondrial matrix Ca^2+^ (**Fig 1D**), closely resembling uptake in response to global KCl global depolarization and resultant Ca^2+^-induced Ca^2+^ release from the ER (**Fig. 1F-G**). This response was specific to Ca^2+^-permeable TPC2 activity, as stimulation with the PI(3,5)P₂ mimetic TPC2-A1-P, which triggers Na^+^ flux through TPC2,^22^ failed to induce mitochondrial Ca^2+^ uptake (**Fig 1E**). Moreover, pretreatment with the TPC2 inhibitor, tetrandrine, abolished the Ca^2+^ signal evoked by TPC2-A1-N (**Fig. 1H-I**), confirming the on-target action of this agonist.

**Figure 1:**
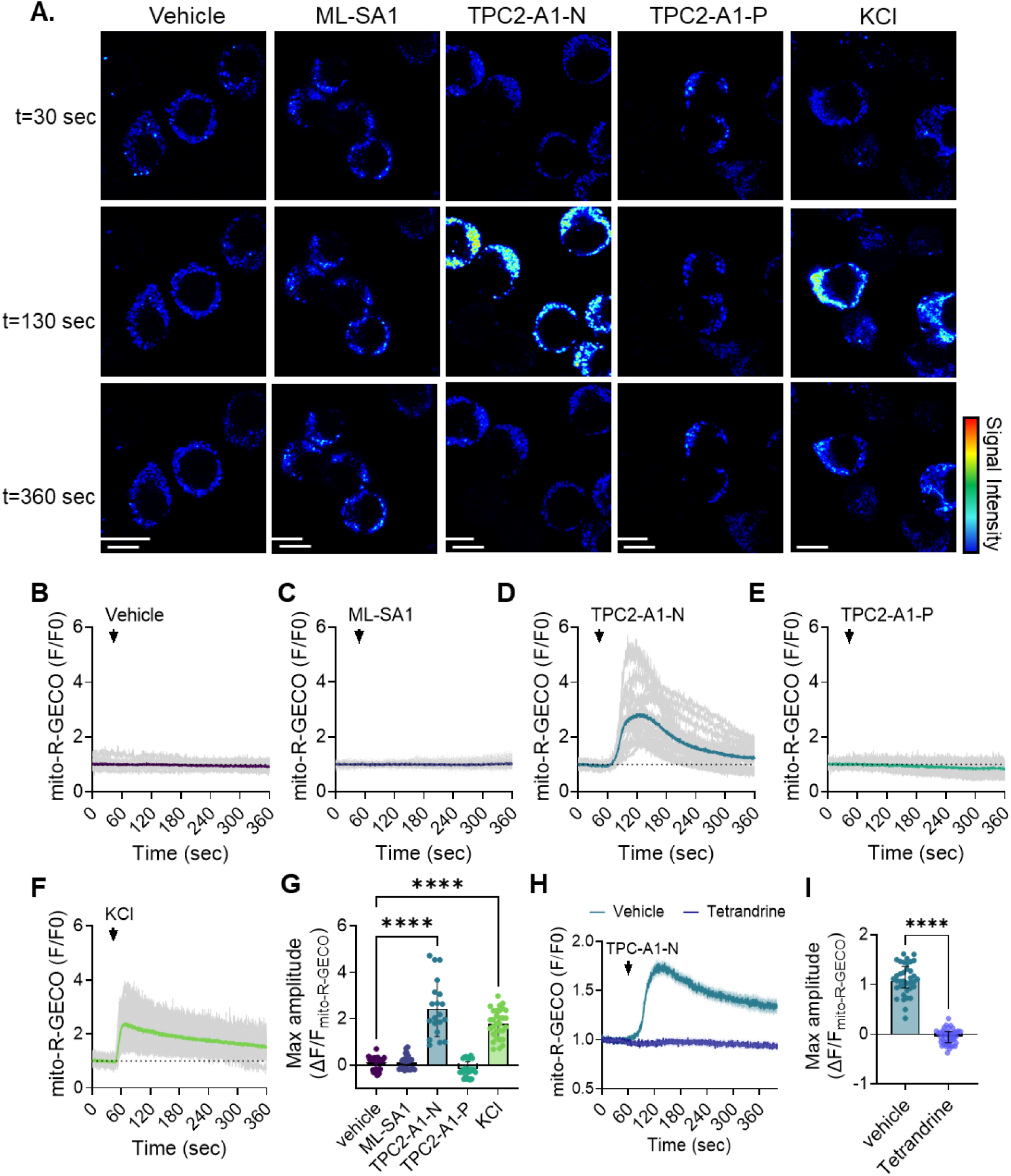
Lysosomal Ca^2+^ release from TPC2 triggers transient mitochondrial Ca^2+^ loading. (**A**) Time-lapse images of mito-R-GECO signal in N2As treated with vehicle, TPC2-A1-N (30 μM), TPC2-A1-P (60 μM), ML-SA1 (30 μM), or KCl (100 mM). Scale bar, 10 µm. (**B-E**) Raw (light gray) and averaged traces showing the change in mito-R-GECO intensity (normalized to baseline) in response to treatment with each lysosomal ion channel agonist or (**F**) KCl. (**G**) Quantification of the maximum peak mito-R-GECO amplitude after treatment. Kruskal-Wallis one-way ANOVA with Dunns post-hoc; \*\*\*\**p*< 0.0001. N=20-37 cells/N=3 days per group. (**H**) Averaged traces showing the normalized mito-R-GECO response to TPC2-A1-N (30 μM) treatment in the presence or absence of tetrandrine (30 μM). (**I**) Quantification of the maximum peak mito-R-GECO amplitude after treatment. Student’s t-test; \*\*\*\**p*< 0.0001. N=37-43 cells/N=3 days per group.

Similar to neurons, human AC16 cardiomyocytes exhibited significant and transient mitochondrial Ca^2+^ uptake in response to TPC2-A1-N, while neither ML-SA1 nor TPC2-A1-P elicited detectable mitochondrial Ca^2+^ signals in these cells (***Suppl. Fig. 1A–D***). To test definitively whether this response was dependent on TPC2, we targeted TPC2 using a CRISPR/Cas9 strategy. Western blotting showed a marked reduction of TPC2 in clonal AC16 cardiomyocytes with no evidence of compensatory upregulation of the related endosomal channel TPC1 (***Suppl. Fig. 1E***). In these TPC2-deficient cells, TPC2-A1-N-evoked mitochondrial Ca^2+^ loading was found to be significantly attenuated, as reflected by reductions in both the peak mito-R-GECO amplitude and total Ca^2+^ load (expressed as area-under-the-curve; ***Suppl. Fig. 1F–H***). Importantly, responses to KCl depolarization remained intact (***Suppl. Fig. 1I-K***), demonstrating that the diminished response reflected specific loss of TPC2 function rather than remodeling of mitochondrial Ca^2+^ uptake pathways. Together, these findings establish TPC2 as a critical mediator of lysosome–mitochondria Ca^2+^ crosstalk in excitable cells.

### TPC2-driven mitochondrial Ca^2+^ uptake requires ER Ca^2+^ release and mitochondrial Ca^2+^ uniporter channel (mtCU) activity

Given the similarity between TPC2- and KCl-induced mitochondrial Ca^2+^ responses, and the known proximity of lysosomes to ER–mitochondrial contact sites,^18,21^ we next tested whether TPC2-mediated mitochondrial Ca^2+^ uptake requires ER Ca^2+^ release via IP_3_ receptors, which are the predominant ER Ca^2+^ release channels in neurons.^1^ For these experiments, N2A cells expressing mito-R-GECO were loaded with the cytosolic Ca^2+^ dye, Fluo4-AM, to simultaneously monitor cytosolic and mitochondrial Ca^2+^ dynamics. TPC2-A1-N was found to evoke a robust increase in cytosolic and mitochondrial Ca^2+^ in N2As, with mirrored amplitudes and kinetics (**Fig 2A,C**). Furthermore, pre-treatment of cells with the IP₃ receptor inhibitor, Xestospongin C, completely abolished both the TPC2-evoked increase in cytosolic and mitochondrial Ca^2+^ (**Fig 2B,D**), indicating that ER Ca^2+^ release is required for the propagation of TPC2-initiated Ca^2+^ signals to mitochondria.

**Figure 2:**
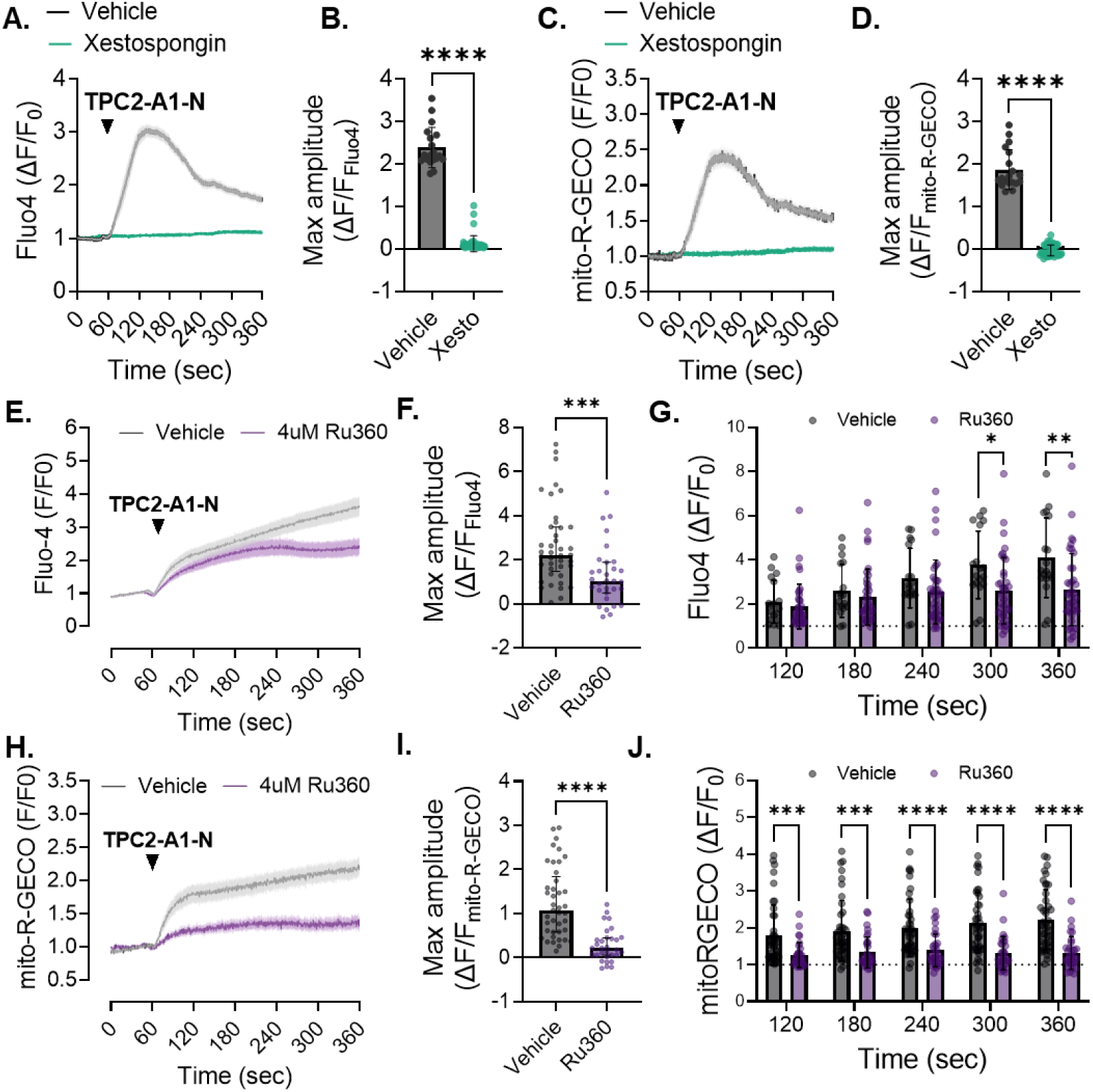
TPC2-dependent Ca^2+^ loading of the mitochondria is ER and MCU-dependent. (**A**) Averaged traces of TPC2-A1-N–evoked cytosolic Ca²⁺ transients in intact N2A cells loaded with Fluo-4 and pre-treated with either vehicle or the IP₃ receptor inhibitor Xestospongin-C (1 μM, 15 min). (**B**) Quantification of peak Fluo-4 amplitudes following TPC2-A1-N stimulation in each treatment group. (**C**) Averaged traces showing simultaneous TPC2-A1-N–evoked mitochondrial Ca²⁺ uptake, measured using mito-R-GECO, in vehicle- and Xestospongin-C–treated N2A cells. (**D**) Quantification of the peak mito-R-GECO amplitude in each group after TPC2-A1-N stimulation. Student’s t-test; \*\*\*\**p*< 0.0001. n=19 (Vehicle) and n=48 (Xestospongin-C) cells/N=3 days per group. (**E**) Averaged traces showing TPC2-A1-N-evoked cytosolic Ca^2+^ transients (Fluo-4), in digitonin-permeabilized N2A cells pre-treated with vehicle or the MCU inhibitor Ru360 (1 μM) prior to stimulation. (**F,G**) Quantification of the maximal (**F**) and time-dependent (**G**) Fluo-4 responses following TPC2-A1-N stimulation in permeabilized cells. (**H**) Averaged mito-R-GECO traces showing TPC2-A1-N–evoked mitochondrial Ca²⁺ uptake in permeabilized cells treated with vehicle or Ru360. (**I,J**) Quantification of the peak (**I**) and time-dependent (**J**) mito-R-GECO responses, demonstrating near-complete suppression of mitochondrial Ca²⁺ uptake by Ru360. Students t-test (F, I) or one-way ANOVA (G, J); \**p* ≤ 0.05, \*\**p* ≤ 0.01, \*\*\**p* ≤ 0.001, \*\*\*\**p* < 0.0001. n=42 (Vehicle) and n=32 (Ru360) cells/N=3 days per group.

We next tested whether mitochondrial Ca^2+^ uptake downstream of TPC2 activation is mtCU-dependent. The mtCU is the reported to be necessary for acute Ca^2+^ uptake across the inner mitochondrial membrane. Since Ru360, a potent inhibitor of the mtCU, displays poor cell penetrability, N2A cells were gently permeabilized with digitonin immediately prior to Ru360 treatment. Under these conditions, TPC2-A1-N stimulation evoked rises in mitochondrial and cytosolic Ca^2+^ levels and co-treatment with Ru360 significantly attenuated both cytosolic and mitochondrial Ca^2+^ responses (**Fig. 2E-J**). Indeed, mitochondrial Ca^2+^ uptake was strongly suppressed across all time points (**Fig. 2H–J**), consistent with a key role for the mtCU in facilitating lysosome-to-mitochondria Ca^2+^ transfer.

### Modulating TPC2 activity tunes the magnitude of mitochondrial Ca^2+^ uptake

Having defined the core components of this Ca^2+^ signaling relay, we next asked whether the magnitude of mitochondrial Ca^2+^ uptake could be amplified by enhancing TPC2 activity. Specifically, we tested whether constitutive enhancement of TPC2 function was sufficient to modulate mitochondrial Ca^2+^ uptake, rather than behaving as an all-or-none mechanism. To address this, we used our N2A TPC2 gain-of-function (GOF) model, in which TPC2 activity is mildly elevated secondarily by genetic ablation of PKARIα-mediated TPC2 inhibition.^23^ Validation of this model is shown in ***Suppl. Fig. 2A–B***, where genetically encoded G-GECO1.2 Ca^2+^ indicators fused to the cytosolic C-terminus of wild-type TPC2, but not the pore-dead mutant (TPC2-D276K), revealed a small but significant increase in baseline TPC2-dependent lysosomal Ca^2+^ efflux, along with an overall increase in agonist-dependent Ca^2+^ release following TPC2-A1-N stimulation (**Fig. 3A–D**). Quite strikingly, TPC2 GOF cells also displayed amplified mitochondrial Ca^2+^ loading in response to TPC2-A1-N, with overall mitochondrial Ca^2+^ content found to be both higher relative to wild-type (WT) N2As (**Fig. 3E–I**). Importantly, KCl-induced mitochondrial Ca^2+^ uptake was comparable between genotypes (***Suppl. Fig. 2C-F***), confirming that the amplified response in TPC2-GOF cells is specific to lysosomal TPC2 activation rather than reflecting altered global Ca^2+^ handling. Consistent with these findings, AC16 TPC2-GOF cardiomyocytes also exhibited amplified mitochondrial Ca^2+^ uptake following TPC2-A1-N stimulation but unaltered responses to KCl depolarization (***Suppl. Fig. 3***), indicating that enhanced lysosome–mitochondria Ca^2+^ coupling by TPC2 is conserved across excitable cell types. Altogether, these findings indicate that TPC2 functions as a tunable Ca^2+^ release channel within the lysosome–ER–mitochondria signaling circuit, capable of magnifying mitochondrial Ca^2+^ load in response to agonist-specific stimulation.

**Figure 3:**
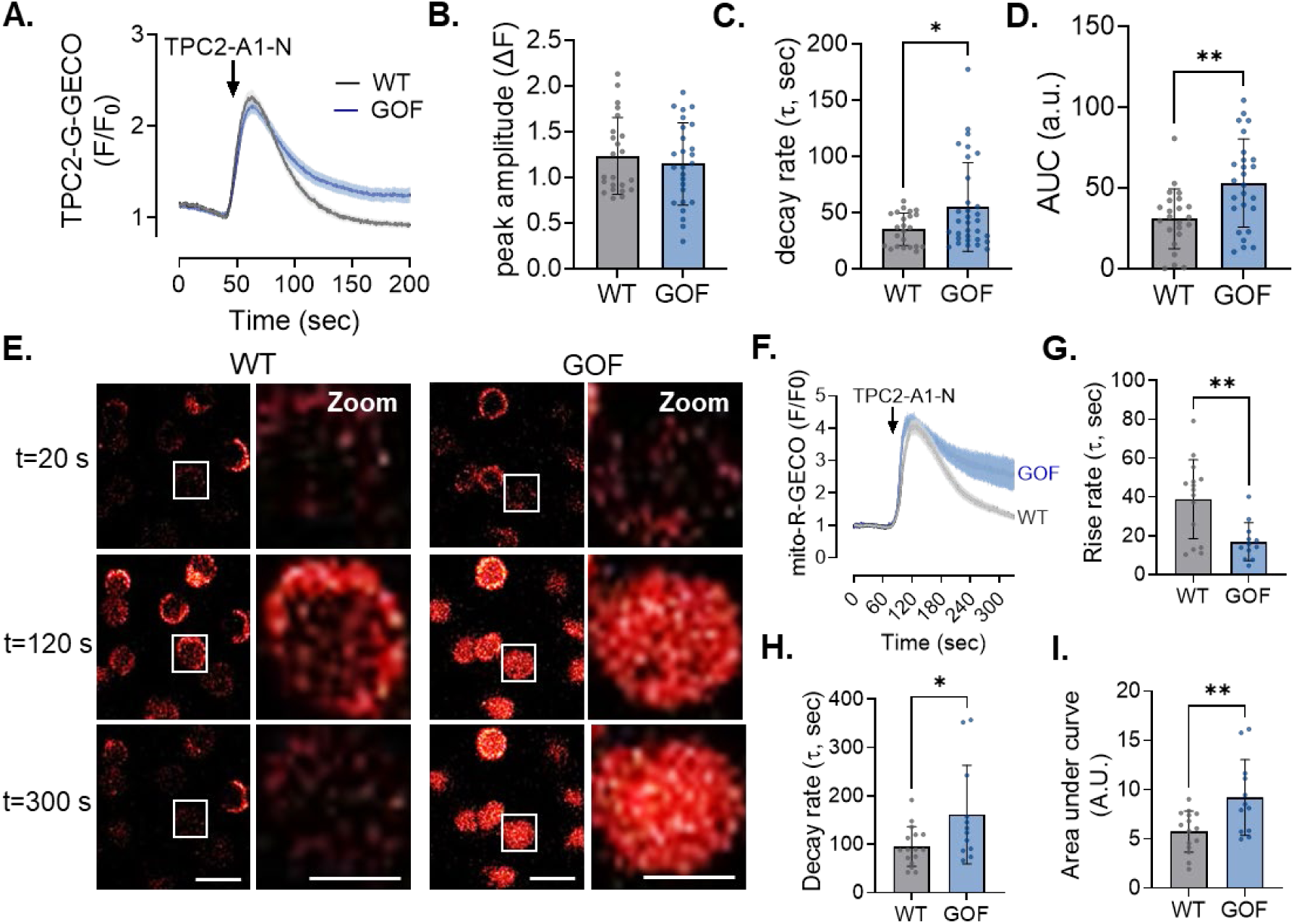
Modulation of TPC2 activity is sufficient to alter mitochondrial Ca^2+^ load. (**A**) Averaged traces of TPC2-A1-N–evoked peri-lysosomal Ca²⁺ transients measured using TPC2-G-GECO1.2 in intact WT and TPC2 GOF N2A cells (n = 23 WT; n = 22 GOF). (**B–D**) Quantification of peri-lysosomal Ca²⁺ responses shown in (A), including peak amplitude (**B**), decay rate (**C**), and area under the curve (**D**). (**E**) Representative time-lapse images of TPC2-A1-N–evoked mitochondrial Ca²⁺ signals (mito-R-GECO) in WT and TPC2 GOF N2A cells, with both wide-field and zoomed views shown (scale bars = 10 µm). (**F**) Averaged traces of TPC2-A1-N–evoked mitochondrial Ca²⁺ transients in WT and TPC2 GOF N2A cells (n = 15 WT; n = 12 GOF). (**G–I**) Quantification of mitochondrial Ca²⁺ dynamics from (F), including rise rate (**G**), decay rate (**H**), and area under the curve (**I**). Student’s t-test; *p ≤ 0.05, **p ≤ 0.01. N=3 independent days per group.

### TPC2 hyperactivation provokes mitochondrial Ca^2+^ overload and increased vulnerability to mPTP opening

One of the best-recognized physiological consequences of an acute increase in mitochondrial Ca^2+^ content is the rapid stimulation of mitochondrial metabolism, driven by an increase in matrix Ca^2+^-stimulated OxPhos and ATP production.^2,24^ As such, we evaluated if lysosomal amplification of mitochondria Ca^2+^ content was sufficient to modulate mitochondrial respiration. For this, we used a Seahorse XF bioanalyzer to measure oxygen consumption rate (OCR) as an index of OxPhos in WT and TPC2-GOF cells. Interestingly, in WT cells acute activation of TPC2 with TPC2-A1-N elicited a transient rise in basal OCR (**Fig. 4A**). ATP-linked respiration and maximal OxPhos capacity remained unchanged (**Fig. 4B**), suggesting that physiological levels of TPC2-driven Ca^2+^ transfer can acutely stimulate mitochondrial respiration without inducing sustained metabolic remodelling. Consistent with this, GOF cells, which exhibit sustained increases in lysosome-to-mitochondria Ca^2+^ flux, displayed elevated basal, maximal, and ATP-linked respiration (**Fig. 4B**, *dark blue bars*). However, upon TPC2-A1-N stimulation, these cells displayed clear evidence of metabolic uncoupling, characterized by an increase in proton leak and a decline in ATP-linked respiration (**Fig. 4B**, *light blue bars*). Use of the TPC2 inhibitor, tetrandrine, confirmed that these genotype-dependent differences in OxPhos arose from chronic TPC2 hyperactivation (**Fig 4C-D**). Importantly, there were no differences in mitochondrial content or electron transport chain (ETC) component expression between genotypes (***Suppl. Fig. 4***), indicating that the observed OxPhos alterations reflect functional changes in mitochondrial activity rather than structural remodeling.

**Figure 4:**
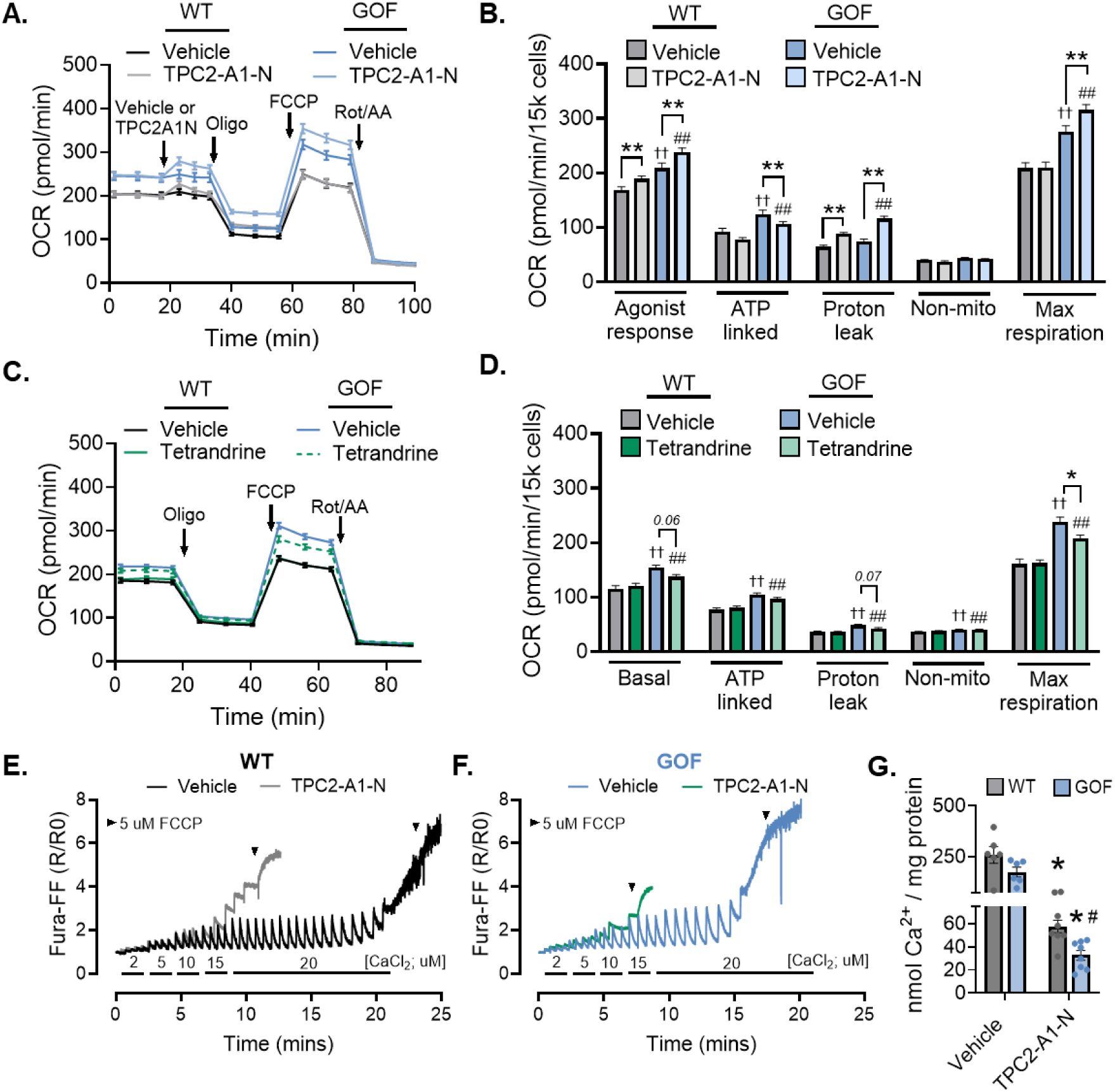
TPC2 hyperactivation provokes mitochondrial Ca^2+^ overload and increased vulnerability to mPTP opening. (**A**) Averaged Seahorse XF traces from WT and TPC2 GOF N2A cells showing oxygen consumption rate (OCR; pmol/min/15,000 cells) after acute treatment with vehicle or TPC2-A1-N (30 µM) following baseline establishment. Sequential injections of oligomycin (Oligo), FCCP, and rotenone/antimycin A (Rot/AA) are indicated. (**B**) Quantification of basal respiration, ATP-linked respiration, proton leak, non-mitochondrial respiration, and maximal respiratory capacity derived from (A). Two-way ANOVA with Tukey post hoc test; **p ≤ 0.01 vs genotype-matched vehicle, ††p ≤ 0.01 vs WT + vehicle, ##p ≤ 0.01 vs WT + TPC2-A1-N. n = 43-49. (**C**) Averaged Seahorse XF traces from WT and TPC2 GOF N2A cells following 20-min pre-incubation with vehicle or tetrandrine (30 µM). (**D**) Quantification of respiratory parameters derived from (C). Two-way ANOVA with Tukey post hoc test; *p ≤ 0.05 vs GOF + vehicle, ††p ≤ 0.01 vs WT + vehicle, ##p ≤ 0.01 vs WT + TPC2-A1-N. n = 50-53. (**E-F**) CRC assay in digitonin-permeabilized WT (**E**) and TPC2 GOF (**F**) N2A cells treated with vehicle or TPC2-A1-N (30 µM) at time 0. Ca²⁺ boluses were added every 60 s until mitochondrial membrane potential collapse (black arrows). Representative traces are shown. (**G**) Quantification of total Ca²⁺ load (nmol/mg protein) tolerated before collapse in the CRC assay. Two-way ANOVA with Tukey post hoc test; *p ≤ 0.05 vs vehicle, #p ≤ 0.05 vs WT. n = 8/group.

Given this divergence between enhanced respiratory drive and metabolic efficiency, we next asked whether sustained TPC2-dependent mitochondrial Ca^2+^ loading increases susceptibility to Ca^2+^-induced mitochondrial permeability transition. Indeed, while moderate elevations in matrix Ca^2+^ stimulate OxPhos, excessive or prolonged mitochondrial Ca^2+^ loading is a well-established trigger of permeability transition pore (mPTP) opening.

To assess mitochondrial susceptibility to Ca^2+^-induced mPTP, we performed Ca^2+^ retention capacity (CRC) assays in permeabilized WT and GOF cells. Briefly, sequential Ca^2+^ boluses were administered while continuously monitoring mitochondrial membrane potential (ΔΨm), and additions were continued until ΔΨm collapse indicated mPTP opening. The cumulative Ca^2+^ load required to trigger depolarization was taken as a measure of mitochondrial CRC. In WT cells, acute treatment with the TPC2 agonist TPC2-A1-N modestly reduced CRC compared to untreated controls (**Fig. 4E**). In contrast, GOF mitochondria displayed a lower basal tolerance to sequential Ca^2+^ pulses, consistent with increased intrinsic sensitivity to Ca^2+^-dependent mPTP opening (**Fig. 4F**). TPC2-A1-N further exacerbated this phenotype in GOF cells, accelerating ΔΨm collapse following fewer Ca^2+^ additions. **Figure 4G** summarizes the quantification of total Ca^2+^ load (nmol/mg protein) tolerated before ΔΨm dissipation across all four groups.

Taken together, these results indicate that TPC2-dependent lysosome–mitochondria Ca^2+^ transfer supports mitochondrial oxidative metabolism under conditions of physiological cell signaling, but that sustained TPC2 hyperactivation accelerates mitochondrial depolarization and Ca^2+^-dependent mPTP opening.

### Modulation of lysosome–mitochondrial Ca^2+^ exchange influences stroke outcomes in vivo

Given our findings that excessive TPC2 activity promotes Ca^2+^-dependent mitochondrial dysfunction, we next examined whether this signaling axis contributes to neuronal injury *in vivo*. We focused on ischemic stroke, a condition in which glutamate-driven Ca^2+^ release and excitotoxicity are major contributors to neuronal death. Because lysosomal TPC2 is a potent amplifier of glutamate-induced intracellular Ca^2+^ release,^25^ we hypothesized that TPC2 hyperactivation would exacerbate ischemic injury by amplifying mitochondrial Ca^2+^ overload during reperfusion.

To test this, we used our TPC2 GOF mouse model^17^ in which TPC2 activity is elevated secondary to genetic ablation of PKARIα-mediated inhibition, mirroring the mechanism used in our cellular studies. WT and TPC2-GOF mice were subjected to transient middle cerebral artery occlusion (tMCAO; 40 mins ischemia followed by 24 hrs reperfusion). Under these conditions, GOF mice displayed markedly poorer survival, with 6 of 9 animals dying within 24 hrs of reperfusion compared with only 2 of 8 WT littermates (**Fig. 5A**). Among surviving animals, infarct size was approximately two-fold larger in GOF mice (**Fig. 5B-C**), accompanied by significantly lower neurological scores at 24 hrs (**Fig. 5C**), indicating worsened functional recovery.

**Figure 5:**
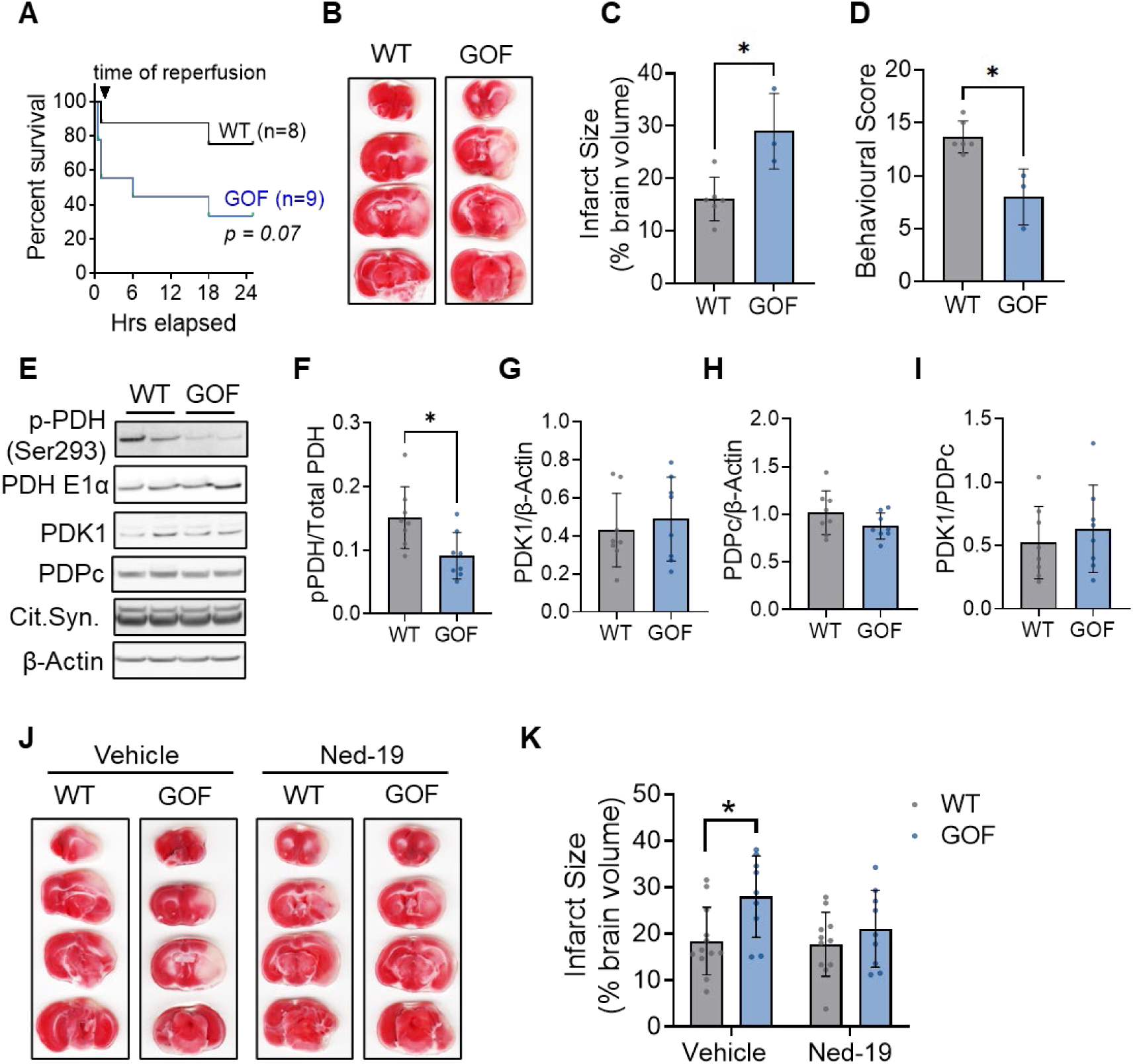
Modulation of lysosome-mitochondrial Ca^2+^ exchange directly influences stroke outcomes in mice. (**A**) Kaplan–Meier survival curves showing overall survival in WT and GOF groups during the first 24 hrs post-tMCAO (40 mins occlusion and 24 hrs reperfusion). (**B**) Representative TTC-stained brain sections in WT and GOF animals from panel A which survived the full 24 hrs post-reperfusion. **(C**) Infarct size calculated from TTC-stained tissue collected 24 hrs post-reperfusion, presented as a percentage of total brain volume. (**D**) Neurological score from the Bederson-Garcia scoring system, carried out blind at 24 hrs post-reperfusion. Mann-Witney U-test; \**p*< 0.05. n=6 (WT) and n=3 (GOF) mice/group. (**E-I**) Western blots (**E**) and quantification of phosphorylated PDH (Ser293), normalized to total PDH E1-α (**F**), PDK1 (**G**), PDPc (**H**), and the PDK1-to-PDPc ratio (**I**) in brains isolated from WT and GOF mice 20 mins post-reperfusion. Relative protein expression was calculated relative to β-actin. Student’s t-test; \**p*< 0.05. n=8/group. (**J**) Representative TTC-stained brain sections in WT and GOF animals at 24 hrs post-tMCAO (30 mins occlusion and 24 hrs reperfusion) treated 5 mins prior to reperfusion with either vehicle or 10μM Ned-19, i.v. (**K**) Infarct size calculated from TTC-stained tissue, presented as a percentage of total brain volume in animals treated with vehicle or Ned-19. Two-way ANOVA with Tukey’s post-hoc analysis; *p<0.05. n=9-12/group.

To assess whether these outcomes were associated with altered mitochondrial Ca^2+^ handling, we next evaluated the phosphorylation status of the Ca^2+^-sensitive matrix enzyme pyruvate dehydrogenase (PDH, Ser-293) in brain lysates collected 20 mins after reperfusion, a time window when mitochondrial Ca^2+^ overload-induced cell death is known to peak.^26,27^ GOF brains exhibited reduced phosphorylation of PDH Ser-293 relative to WT (**Fig. 5E-F**), consistent with enhanced mitochondrial Ca^2+^ uptake. This was not accompanied by genotype-dependent differences in PDK1 or PDPc expression, or in the PDK1/PDPc ratio (**Fig. 5G-I**), indicating that reduced phospho-PDH reflected elevated matrix Ca^2+^ rather than altered regulatory enzyme abundance.

Building on these findings, we next asked whether acute pharmacological inhibition of TPC2 could mitigate the heightened ischemic vulnerability observed in GOF mice. To improve survival, a less severe tMCAO protocol (30 min ischemia, 24 hrs reperfusion) was used, under which all animals survived. Vehicle-treated GOF mice continued to exhibit significantly larger infarcts compared with WT controls (**Fig. 5J**). Strikingly, intravenous administration of the TPC2 inhibitor Ned-19 (10 µM) 5 mins prior to reperfusion prevented this genotype-dependent increase in infarct size, normalizing infarct volume in GOF mice to WT levels (**Fig 5K**).

### Pharmacological inhibition of TPC2 confers neuroprotection in human models of stroke injury

Finally, to determine whether pharmacological TPC2 inhibition can confer neuroprotection in a translational context, we next used human induced pluripotent stem cell (iPSC)–derived neurons subjected to an *in vitro* model of I/R (**Fig 6A**). Neurons were exposed to oxygen–glucose deprivation (OGD) followed by reoxygenation, which recapitulates the metabolic and excitotoxic stress seen during cerebral reperfusion. Under these conditions, neuronal death increased in both an OGD duration-dependent and reperfusion-dependent manner (**Fig. 6B-D**), confirming robust induction of I/R injury.

**Figure 6:**
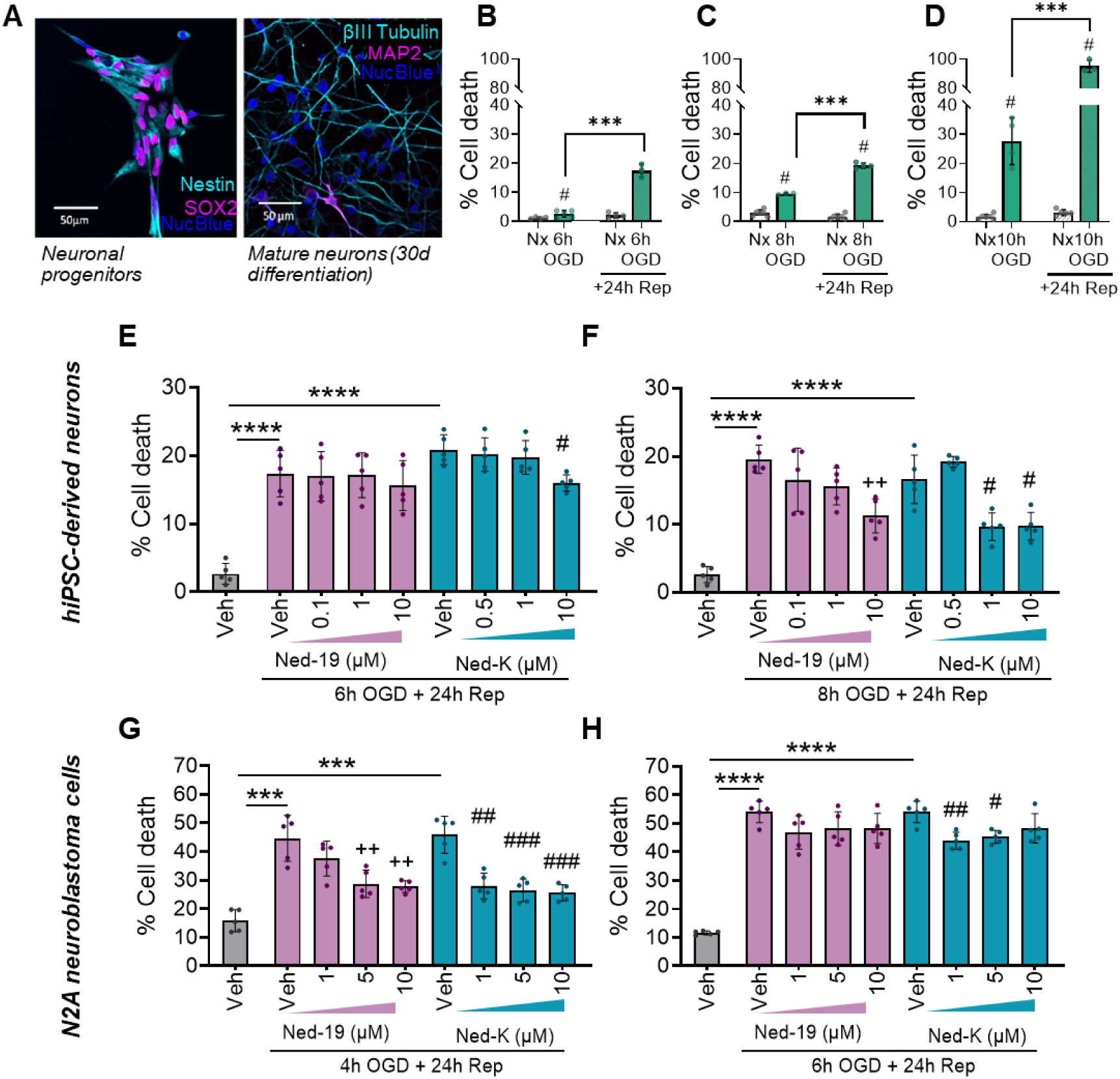
Pharmacological TPC2 inhibition provides potent neuroprotection in *in vitro* models of stroke. (**A**) Representative images of Nestin+, SOX2+ neuronal progenitor cells and mature neurons (identified using βIII-Tubulin and MAP2) after 30 days of differentiation. Nuclei stained using Hoechst 33342. (**B-D**) Percentage cell death in mature human iPSC-derived neurons in response to (**B**) 6 hrs, (**C**) 8 hrs or (**D**) 10 hrs of oxygen-glucose deprivation (OGD) alone (solid bars) or OGD with 24 hrs of reperfusion (dashed bars) which, as expected, induces substantially more cell death. (**E**) Percentage cell death in human iPSC-derived neurons in response to 6 hrs of OGD and 24 hrs of reperfusion in the presence of increasing doses of Ned-19 or Ned-K. (**F**) As E but 8 hrs of OGD. (**G**) Percentage cell death in mouse N2A cells in response to 4 hrs of OGD and 24 hrs of reperfusion in the presence of increasing doses of Ned-19 or Ned-K. (**H**) As in G but 6 hrs of OGD. Nx = normoxia, Veh. = vehicle treatment in the absence (grey bars) or presence (colored bars) of OGD. B-D analyzed using a two-way ANOVA and E-H analysed using a repeated measures two-way ANOVA with Tukey post-hoc analysis. ***p ≤ 0.001, ††p ≤ 0.01 vs Vehicle for Ned-19 treated group, #p ≤ 0.05 vs Vehicle for Ned-K treated group. Panels B-C, n=3-6/group. Panels E=H, n=5/group.

We next examined whether pharmacological inhibition of TPC2 could limit neuronal injury following reperfusion. For this, human iPSC-derived neurons and N2A cells were both treated with the TPC2 inhibitor Ned-19 at the time of reoxygenation, and cell death was quantified 24 hrs later. Ned-19 significantly reduced cell death in a dose-dependent manner in both neuronal models compared with vehicle-treated controls (**Fig. 6F-G**). Because Ned-19 has moderate potency and limited selectivity, we next evaluated Ned-K, a next-generation TPC2 inhibitor with improved binding affinity.^28^ In human iPSC-derived neurons, Ned-K administration at the time of reoxygenation significantly reduced neuronal death at lower concentrations, as compared with Ned-19 (**Fig. 6E-F**). Parallel experiments in N2A cells subjected to the same OGD/reoxygenation paradigm yielded comparable protective effects (**Fig. 6G-H**), indicating that enhanced pharmacological inhibition of TPC2 robustly limits reperfusion-induced neuronal death across models.

Collectively, these results demonstrate that acute pharmacological inhibition of TPC2 confers dose-dependent neuroprotection in both murine and human neuronal models of I/R injury. Moreover, they establish that more potent TPC2 inhibitors provide substantial efficacy, reinforcing lysosomal TPC2 as a therapeutically tractable target for limiting reperfusion-induced neuronal injury.

## Discussion

In this study, we define a lysosome-ER-mitochondria signaling pathway in which TPC2-dependent lysosomal Ca^2+^ release governs mitochondrial bioenergetic output and stress susceptibility. We show that TPC2 activation initiates ER-dependent Ca^2+^ relay through IP₃ receptor engagement, resulting in enhanced mitochondrial Ca^2+^ uptake via mtCU. The magnitude and duration of mitochondrial Ca^2+^ accumulation scale with TPC2 activity, such that moderate activation transiently enhances OxPhos, whereas sustained or excessive activation promotes mitochondrial depolarization, vulnerability to mPTP, and exacerbates cell death during ischemic stress. In vivo, modulation of TPC2 activity directly alters mitochondrial Ca^2+^ load and infarct severity following transient I/R. We propose a model in which TPC2 functions as a regulatory amplifier within confined lysosome-ER-mitochondria nanodomains, tuning mitochondrial energetics under basal conditions but predisposing cells to Ca^2+^ overload and injury when hyperactivated.

Past studies have positioned lysosomes as either passive participants in mitochondrial quality control (e.g., via participation in mitophagy)^29^ or as indirect regulators of mitochondrial homeostasis through TRPML1-dependent mitochondrial fission.^13,18^ Our data instead support a model in which lysosomal TPC2 activity directly regulates mitochondrial bioenergetics through spatially confined Ca^2+^ relay across lysosome-ER-mitochondria junctions. We find that TPC2 activation increases mitochondrial matrix Ca^2+^ content in proportion to channel activity, yet without commensurate changes in bulk cytosolic Ca^2+^ transients. This dissociation between local mitochondrial Ca^2+^ accumulation and global Ca^2+^ handling argues that TPC2-dependent signaling operates within restricted inter-organelle nanodomains rather than through diffuse cytosolic amplification. Notably, Ca^2+^ concentrations within the TPC2 nanodomain are estimated to reach levels up to two orders of magnitude greater than global cytosolic Ca^2+^,^30^ providing a quantitative basis for how spatially confined lysosomal Ca^2+^ release can exert potent downstream effects despite minimal changes in whole-cell Ca^2+^ signals. This interpretation is consistent with prior observations that TPC2 activation can evoke highly localized ER Ca^2+^ release events (“tuffs”) without substantially altering whole-cell Ca^2+^ dynamics.^31^ Similarly, in our previously characterized TPC2 GOF cardiomyocytes, lysosomal Ca^2+^ release triggered spontaneous ER/SR Ca^2+^ release without modifying Ca^2+^-induced Ca²⁺ release or global Ca^2+^ cycling.^17^ As such, we propose a hierarchical model in which lysosomal Ca^2+^ release serves as an initiating signal that is selectively amplified at ER–mitochondria contact sites. Although speculative, it’s feasible that such amplification occurs at mitochondria-associated membranes (MAMs), where close apposition of ER and mitochondrial membranes enables high-fidelity Ca^2+^ transfer independent of bulk cytosolic diffusion.^32^ In this regard, TPC2 would function as an adjustable gain control upstream of IP₃R- and mtCU-dependent transfer, scaling the Ca^2+^ load delivered to mitochondria without globally perturbing cytosolic Ca^2+^ homeostasis. Such spatial compartmentalization provides a mechanistic explanation for how relatively modest alterations in lysosomal channel activity can produce disproportionate effects on mitochondrial energetics and cell fate.

Our findings add further to the growing evidence that, despite residing on the same organelle, TRPML1 and TPC2 generate autonomous Ca^2+^ nanodomains which are functionally non-redundant. Although few studies have directly compared the physiological roles of these lysosomal Ca^2+^ channel families, available data consistently demonstrate selective downstream coupling to either TPC2^22,33,34^ or TRPML1,^35,36^ supporting the idea that they operate as independent Ca^2+^ efflux pathways. This functional segregation has been directly demonstrated by Davis et al., who used channel-fused genetically encoded Ca^2+^ indicators – including the same G-GECO constructs used here – to quantify channel-specific nanodomain Ca^2+^ signals, revealing that TPC2- and TRPML1-derived Ca^2+^ nanodomains are largely insulated from one another.^30^ This insulation provides a mechanistic explanation for why Ca^2+^ efflux through these channels drives distinct physiological outcomes that cannot be compensated for by the other.^33^ Importantly, this segregation cannot be explained solely by differences in ligand activation. While TRPML1 is activated by the lysosome-specific lipid PI(3,5)P₂ and TPC2 by the cytosolic second messenger NAADP,^37–39^ recent optical imaging studies have demonstrated cooperative enhancement of TPC2-mediated Ca^2+^ release by PI(3,5)P₂,^31^ indicating that shared lipid signaling can stimulate Ca^2+^ efflux through both channels under appropriate conditions. An alternative hypothesis is that each channel is preferentially coupled to distinct Ca^2+^ decoding machinery within its local nanodomain, thereby conferring signaling specificity. However, both our findings and those of Peng et al.^13^ demonstrate that mitochondrial Ca^2+^ uptake downstream of TPC2 and TRPML1, respectively, requires mtCU activity yet regulate distinct physiological functions, suggesting convergence on a common Ca^2+^ uptake pathway rather than exclusive coupling to distinct decoders. Instead, functional specificity may arise from spatial or molecular heterogeneity among lysosomal subpopulations, including differences in TPC2 and TRPML1 expression, distribution, or stoichiometry, a possibility that warrants further investigation.

We observe that mitochondrial responses to TPC2 activation are nonlinear, consistent with a threshold-dependent Ca^2+^ regulatory system. Specifically, moderate TPC2 activation acutely enhances OxPhos without causing depolarization. Based on well-established principles, we infer that this effect likely reflects stimulation of pyruvate dehydrogenase, TCA cycle flux, and ATP synthesis via mitochondrial matrix Ca^2+^-sensitive enzymes, linking energy production to cellular demand.^2,24,40,41^ In contrast, sustained or excessive TPC2 activation triggers rapid and irreversible mitochondrial depolarization and permeability transition, with concomitant cell death. These findings indicate that TPC2 defines both the amplitude and duration of Ca^2+^ exposure within a narrow functional window that separates metabolic adaptation from injury. In this framework, lysosomal TPC2 serves as a graded upstream control point, determining whether mtCU-dependent Ca^2+^ entry remains bioenergetically supportive or becomes pathological.

This threshold behavior is particularly relevant in I/R injury, where mitochondrial Ca^2+^ overload is an early and sustained driver of neuronal damage.^27,42,43^ While canonical models emphasize glutamate-driven ER Ca^2+^ release as the principal driver of neuronal excitotoxicity, prior studies demonstrate that glutamate also stimulates NAADP production, activating TPC2-dependent lysosomal Ca^2+^ release and positioning lysosomes as potent amplifiers of excitotoxic mitochondrial Ca^2+^ accumulation.^25,44^ Our findings extend this paradigm by showing that TPC2 activity quantitatively governs mitochondrial matrix Ca^2+^ accumulation and permeability transition during ischemic stress in vivo. Indeed, in TPC2 GOF mice even modest enhancement of channel activity was sufficient to exacerbate infarct size, worsen neurological outcomes, and increase mortality following transient middle cerebral artery occlusion. Biochemical evidence of reduced PDH phosphorylation during early reperfusion further supports enhanced mitochondrial Ca^2+^ loading *in vivo*. Notably, acute pharmacological inhibition of TPC2 at the onset of reperfusion normalized infarct burden, demonstrating that this pathway is not merely correlative but functionally involved in determining injury severity.

The translational implications of these findings are strengthened by our observations in human iPSC-derived neurons. Pharmacological inhibition of TPC2 at the time of reoxygenation conferred dose-dependent neuroprotection in both murine and human neuronal models of I/R injury, with next-generation inhibitors demonstrating enhanced potency at lower concentrations. These results suggest that targeting lysosomal Ca^2+^ amplification during the narrow therapeutic window of reperfusion may limit mitochondrial Ca^2+^ overload while preserving basal metabolic signaling. Importantly, such an approach does not require complete suppression of mitochondrial Ca^2+^ uptake, potentially avoiding the metabolic liabilities associated with direct mtCU inhibition.^45–47^

While our findings establish a functional lysosome–mitochondrial signaling axis that shapes cellular stress adaptation, additional mechanistic questions remain to be resolved. We did not directly visualize Ca^2+^ nanodomains at lysosome–ER–mitochondria interfaces, nor determine whether ischemic stress dynamically remodels organelle contact architecture, which may influence the efficiency of Ca^2+^ relay under pathological conditions. Additionally, while TPC2 hyperactivation increases the vulnerability to mPTP opening, the molecular coupling between lysosomal Ca^2+^ flux and permeability transition warrants further investigation. Nonetheless, our data firmly establish functional causality between lysosomal TPC2 activity, mitochondrial Ca^2+^ loading, and tissue injury with clear evidence of therapeutic potential.

In summary, we identify lysosomal TPC2 as an amplifier of a multi-organelle Ca^2+^ signaling network that dictates mitochondrial energetic output and cell death. These findings expand the role of lysosomes in cellular stress responses and reveal inter-organelle Ca^2+^ transfer as a regulated system, rather than a binary or independent event. Therapeutic modulation of lysosomal TPC2-dependent calcium flux may therefore represent a strategy to recalibrate mitochondrial vulnerability in acute ischemic injury.

## Methods

### Mouse neuronal and human cardiomyocyte cell lines

Mouse neuroblastoma Neuro-2A (N2A; ATCC CCL-131) cells were maintained in DMEM supplemented with 10% fetal bovine serum (FBS) and 1% penicillin/streptomycin. Human cardiomyocyte AC16 cells (Sigma #SCC109) were cultured in DMEM/F12 (1:1) supplemented with 12.5% FBS and 1% penicillin/streptomycin. All cells were maintained at 37 °C in a humidified incubator with 5% CO₂ and routinely tested for mycoplasma contamination. As AC16 human cardiomyocytes are known to be phenotypically altered with extended passage,^48^ all experiments using AC16s were performed using cells with passage numbers <12.

A TPC2 gain-of-function (GOF) N2A cell line was generated by introducing a single-nucleotide substitution (TGC→AGC; C18S) into the *prkar1a* gene, which we previously demonstrated increases TPC2 channel activity by relieving an inhibitory regulatory interaction.^17^ CRISPR-Cas9–mediated genome editing was performed by Thermo Fisher Scientific using their standard in-house protocols. An in vitro–transcribed guide RNA (gRNA; 5′-GCACATAGAGCTCGCATTCC-3′) and a symmetric single-stranded oligonucleotide donor (sense strand) were designed to target the desired locus and co-electroporated into N2A cells together with Cas9 nuclease. Editing efficiency was assessed by next-generation sequencing, after which the edited pool was subjected to limiting dilution cloning. Individual colonies were expanded and screened by Sanger sequencing to identify clones homozygous for the C18S mutation.

TPC2 knockout (TPC2 KO) AC16 cells were generated using CRISPR-Cas9 technology. A custom plasmid (VectorBuilder) expressing Cas9(D10A) under the CBh promoter, dual gRNAs targeting human TPCN2 (chromosome 11q13.3) under U6 promoters (gRNA#6196, 5′-ACTGGTACTCGGGCCTCGGC-3′; gRNA#6213, 5′-CCTCATCGCCCTGGCAAACC-3′), and a blasticidin resistance cassette under a CMV promoter was used. AC16 cells were transfected with jetPRIME according to the manufacturer’s protocol and selected with blasticidin (10 µg/mL) until all untransfected control cells were eliminated. Surviving cells were diluted and plated into 15 cm dishes for clonal isolation using glass cloning cylinders. Knockout efficiency was validated by quantitative PCR and Western blot analysis.

### Live cell Ca^2+^ transient imaging

For Mito-R-GECO experiments, N2a WT and TPC2 GOF cells were transduced with adenovirus encoding the mitochondrial Ca^2+^ reporter Mito-R-GECO 48 hrs before imaging.^49^ Immediately before imaging, cells were washed once in Tyrode’s buffer solution (130 mM NaCl, 5.6 mM KCl, 3.5 mM MgCl2ꞏ6H2O, 17.5 mM glucose, 5 mM HEPES, 1.4 mM CaCl2ꞏ2H2O, pH = 7.4) and, where cytosolic Ca^2+^ was simultaneously obtained, loaded with Ca^2+^ sensitive dye Fluo-4-AM at a final concentration of 2 ug/ml. Cells were imaged in fresh Tyrode’s buffer in a 37°C chamber using a Carl Zeiss Axio Observer Z1 microscope. Cytosolic and/or mitochondrial Ca^2+^ signals were continuously recorded for 6-8 mins at 490/20 nm_ex_, 490/20 nm_ex_-632/60 nm_em_, and 572/35 nm_ex_-632/60 nm_em_ respectively. After 60 s of baseline recording, a bolus of KCl (100 mM), ML-SA1 (30 μM), TPC2-A1-N (30 μM), TPC2-A1-P (60 μM) or vehicle (DMSO) was added. For experiments with Xestosponsgin-C treatment to block IP3 receptors, cells were incubated in 1 µM Xestosponsgin-C prepared in Tyrode’s buffer solution for 15 min before imaging. To permeabilize cells for Ru360 (4 μM) treatment, Mito-R-GECO transduced cells were washed with a permeabilization buffer (120 mM KCl, 10 mM NaCl, 1 mM KH_2_PO_4_, 20 mM HEPES, 5 mM succinic acid, protease phosphatase inhibitor, 80 μg/ml digitonin, 0.2 μM CaCl_2_) and then imaged as above.

For experiments with TPC2-G-GECO and D276K-TPC2-G-GECO plasmids (a gift from Antony Galione,^33^ Addgene plasmid #207141 and 207142, respectively), N2a WT and TPC2 GOF cells were transiently transfected with the plasmids using Jetprime transfection reagent (VWR # 89129-924) and allowed to express for 48 hrs before imaging. For imaging, cell baselines were recorded for 60 s, after which a bolus of TPC2-A1-N (30 μM) was added. Ca^2+^ signals were continuously recorded for 6 mins at the EGFP channel at 490/20 nm_ex_, 632/60 nm_em_. Ca^2+^ transients were analyzed in Clampfit 10.7 software.

### Oxygen Consumption Rate Assays

A Seahorse Bioscience XF96 extracellular flux analyzer was employed to measure oxygen consumption rates (OCR) in N2A cells. 15,000 N2A WT and TPC2 GOF cells per well were plated in DMEM base media, pH 7.4, supplemented with 1 mM pyruvate and 2 mM L-glutamine. After measuring basal OCR, TPC2 agonist TPC2-A1-N (30 μM) or vehicle was injected, followed by 3 μM oligomycin injection to inhibit ATP-linked respiration, 3 μM FCCP to measure maximal respiration, and finally to completely inhibit all mitochondrial respiration, 1 μM rotenone/10 μM antimycin-A was injected. For subsequent experiments evaluating the contribution of sustained TPC2 hyperactivation on increased OCR in GOF cells cells were pre-incubated with either vehicle or the TPC2 inhibitor, tetrandrine (30 μM), 20 minutes prior to the OCR assay.

### Mitochondrial Ca^2+^ retention assay

To perform mitochondrial Ca^2+^ retention capacity (mCRC) experiments N2A cells were transferred to an intracellular-like medium containing 120 mM KCl, 10 mM NaCl, 1 mM KH2PO4, 20 mM HEPES-Tris, and 3 μM thapsigargin to inhibit SERCA so that the movement of Ca^2+^ was only influenced by mitochondrial uptake, 40 μg/mL digitonin, protease inhibitors (Sigma, EGTA-Free Cocktail), supplemented with 10 μM succinate and pH to 7.2. All solutions were cleared with Chelex 100 to remove trace Ca^2+^ and experiments were conducted at 37 °C. 2 x 10^6^ digitonin-permeabilized N2A cells were used for simultaneous ratiometric monitoring of bath Ca^2+^ levels (Fura-FF) and Ψm (JC-1) as cells were exposed to consecutive boluses of Ca^2+^ at increasing concentration (2-20 mM Ca^2+^) until Ψm collapsed indicating membrane permeability transition. At completion of the experiment 10 μM FCCP was added to uncouple the ΔΨm and release matrix free-Ca^2+^. Cells were then lysed to quantify protein and the total amount of Ca^2+^ that cells could take up before Ψm collapsed was calculated and expressed as nmol Ca^2+^/mg protein.

### Animal studies

Male and female mice, 12–18 weeks of age, were housed under standard diurnal lighting conditions (12 hr light/dark cycle) with ad libitum access to food and water. All procedures were carried out in accordance with the UK Animals (Scientific Procedures) Act (1986) and were approved by local ethical review panels (LERP and ACER, University of Oxford) under license number P6E80C21D. All experiments adhered to ARRIVE and IMPROVE (stroke-specific) guidelines. The TPC2 GOF model used corresponds to a knock-in mouse line on a C57BL/6 background harboring a mutation in the regulatory subunit Iα of protein kinase A (PKARIα) that results in constitutively enhanced TPC2 activity.^17^ WT littermates were used as controls in all experiments.

### Transient Middle Cerebral Artery Occlusion (tMCAO)

Animals were randomly assigned to treatment groups, and all surgeries, treatments, outcome assessments, and data analyses related to tMCAO experiments were performed with investigators blinded to genotype and treatment group.

Mice were anesthetized with 2% isoflurane in a 60:40 oxygen:nitrous oxide mixture (1 L min⁻¹). Following a midline cervical incision, the common carotid artery (CCA) was isolated and temporarily ligated. The external carotid artery (ECA) was isolated and permanently ligated by electrocautery, and the internal carotid artery (ICA) was isolated and transiently occluded using a microvascular clamp. An arteriotomy was made in the ECA, and a silicone-coated monofilament (6/0, 60L12C; Doccol, USA) was introduced and advanced through the ICA to occlude the origin of the middle cerebral artery at the circle of Willis. The filament was secured in place and maintained for either 30 or 40 min, as specified in the Results. Reperfusion was initiated by gentle withdrawal of the filament, after which incisions were closed and animals were allowed to recover for 24 hrs. For pharmacological studies, a 31-gauge tail vein cannula was inserted 5 min prior to reperfusion for intravenous administration of Ned-19 (10 µM) or vehicle. In experiments requiring tissue collection at 20 min post-reperfusion, mice were maintained under anesthesia throughout the reperfusion period, after which brains were rapidly harvested and snap-frozen.

### Behavioural Analysis

Neurological deficits were assessed using a modified Bederson–Garcia scoring system taken from^50^ which is commonly used to evaluate motor impairment following experimental stroke in mice. Briefly, mice were evaluated 1 day prior to, and at 24 hrs post-stroke on the Bederson scale^51^ for flexion (0-1), response to lateral push (0-1) and circling (0-1) and on the Garcia scale^52^ for spontaneous activity (0-3), four limb symmetry (0-3), forepaw outstretching (0-3), climbing (0-3), body proprioception (0-3) and vibrissae touch (0-3). This modified scale should give a score of 20 for a totally healthy animal.

### Infarct assessment

Animals were euthanized by cervical dislocation and transcardially perfused with heparinized saline. Brains were rapidly removed and sectioned into six 1-mm thick coronal slices using a brain matrix. Sections were incubated in 1% triphenyltetrazolium chloride (TTC) at 37 °C for 30 min to differentiate viable tissue (TTC-positive) from infarcted tissue (TTC-negative). Sections were scanned, and infarct area was quantified using ImageJ and expressed as a percentage of total brain volume.

### Human iPSC-derived neurons

HPSI0114i-kolf_2-C1 cells generated from KOLFC2 cells by the Wellcome Trust Sanger Institute were transformed into neural progenitor cells (NPCs) via small molecule dual SMAD inhibition using 10 μM SB431542 (Cambridge Bioscience; #ZRD-SB-50) and 100nM LDN193189 (Stemcell Technologies; # 72147) in place of dorsomorphin. NPCs were maintained in culture with ENSTEM neural medium with 2mM L-Glutamine and 20ng/ml fibroblast growth factor-basic, changed on alternate days until cells reached a density of 0.5x10cm^-2^ at which time, cells were split and differentiated to mature neurons. For differentiation, NPCs were seeded in 6 well plates at a density of 1x10^5^ cells/cm^2^ with forebrain differentiation basal media for 6 days. At this point, cells were detached using Accutase^TM^ and seeded at 2-4 x10^4^cm^-2^ in Brain Phys Neuronal Medium supplemented with STEMdiff^TM^ Forebrain Neuron Maturation Supplement. Media was changed every 3 days and cells matured over 30 days.

Characterization of NPCs and 30-day differentiated KOLF hiPSCs was done by immunostaining using the NPC markers Nestin (mouse anti-human, 1:250; [R&D systems; #656801]), SOX2 (goat anti-human SOX2, 1:100; [R&D systems #AF2018-SP]), the mature neuronal marker microtubule-associated protein 2 (MAP2) (rabbit anti-human MAP2, 1:200; [Genetex GTX133109]), and β-III tubulin (mouse anti-human β-III tubulin, 1:500; [DSHB #E7]). For all staining, cells were first fixed for 15 mins with 4% PFA, washed 3x using PBS-T, and then permeabilized for 10 minutes using PBS-T with 0.1% Triton. Cells were blocked for 30 minutes using blocking buffer (PBS-T, 10% donkey serum, 0.1% BSA) and then stained for 1 hour at room temperature with primary antibodies diluted in blocking buffer. Secondary antibodies against mouse (AF-488), rabbit (AF-488 and AF-555), or goat (AF-488 and AF-555) were applied, as appropriate. Cells were washed 2x with PBS-T before Hoechst nuclear staining (1:2000) was applied. Cells were mounted using ProLong^TM^ Diamond Antifade (Thermo Fisher Scientific) before images were acquired with a Leica TCS SP5 X confocal microscope equipped with a 63X oil immersion lens.

### In vitro model of stroke

Oxygen–glucose deprivation (OGD) was used to model ischemic stroke in vitro using a hypoxia workstation. Glucose-free culture medium and phosphate-buffered saline (PBS) were pre-equilibrated overnight in the hypoxia chamber (0% O₂). Human iPSC-derived neurons were then transferred into the workstation, and culture medium was replaced with OGD medium for the indicated durations. For OGD-only conditions, cells were lysed immediately at the end of the hypoxic period. To model reperfusion, cultures were removed from the hypoxia chamber, OGD medium was replaced with standard glucose-containing medium, and cells were returned to normoxic conditions (21% O₂) for 24 hrs prior to analysis.

### Cell death assays

The percentage of cell death following OGD or OGD with reperfusion was quantified using the CytoTox 96® non-radioactive cytotoxicity assay (Promega, G1780), according to the manufacturer’s instructions. Lactate dehydrogenase (LDH) release was measured from experimental wells and normalized to a maximal lysis control, generated by adding lysis buffer to untreated wells seeded at identical cell density. Percent cell death was calculated relative to this maximal LDH release after subtraction of background signal from media-only controls. To assess the contribution of lysosomal Ca^2+^ signaling to neuronal cell death following OGD/reperfusion, cells were treated with increasing concentrations of the TPC2 inhibitors *trans-*Ned-19, Ned-K, or vehicle control (0.1% DMSO final concentration), as indicated.

### Immunoblotting

For PDH A1 and phospho PDH A1 (Ser293) immunoblotting, snap-frozen brain tissue was lysed in 1x RIPA buffer (Cell signaling # 9806) with 1x protease phosphatase inhibitor cocktail (Thermo # A32961), bead homogenized and then spin-clarified for 10 minutes at 14000 rpm to obtain protein samples. Following the addition of 1x LDS buffer (Thermo # NP0007) mixed with 1x reducing buffer (Thermo # NP0009) and boiling at 95 ^0^C for 10 minutes, the lysate was loaded onto a 10% Tris-Glycine gel and subsequently transferred to the nitrocellulose membrane using a constant current of 250 mA. Membranes were blocked in Rockland blocking buffer (Thomas # MB-070) and then switched to primary antibody (rabbit anti-phospho PDH A1 (Ser293) [Abcam #ab92696], 1:1000 in Rockland blocking buffer) and secondary antibody (IRDye 800CW goat anti-rabbit [LI-COR # 92668071], 1:10,000). After imaging in the LICOR imaging system, the membrane was stripped (Thermo # 46430), washed in 1x TBST, blocked, and subsequently incubated in mouse anti-PDH A1 (Abcam # ab110330, 1:1000) and secondary antibody (IRDye 800CW goat anti-mouse [LI-COR # 92632210], 1:10,000) diluted in Rockland buffer. Band intensity of total PDH and phospho PDH was quantified in ImageJ software to obtain their ratio. Rabbit anti-citrate synthase (Cell signaling #14309, 1:1000), mouse anti-PDK1 (Santa Cruz #sc-515944, 1:1000), mouse anti-PDPc (Santa Cruz #sc-398117, 1:1000), total OxPHOS rodent WB antibody cocktail (Abcam #ab110413, 1:1000) were used in other experiments. For TPC2 KO validation, AC16 cardiomyocyte cells were lysed in solubilization buffer (150 mM NaCl, 20 mM HEPES, 1% w/v CHAPS, and 1x protease phosphatase inhibitor cocktail, pH=7.2), and reduced in 1x LDS buffer mixed with 100 mM DTT. Rabbit anti-TPC2 (Sigma #HPA027080, 1:500) and donkey anti-rabbit-HRP (Cell signaling #7074) antibodies were used as primary and secondary antibodies respectively.

### Statistics

Data were checked for normality of distribution before statistical analysis using Shapiro-Wilk’s normality test. Comparisons between normally distributed data were performed using either a Student’s t-test or an analysis of variance (ANOVA) with Tukey’s correction, as indicated in the figures. Non-normally distributed data were compared using the Mann-Whitney test or Kruskal-Wallis test. Results were considered statistically significant at a p-value <0.05.

## Data availability

All data supporting the findings of the study are available from the corresponding author upon reasonable request.

## Acknowledgements

This work was supported by NIH grant R01 HL180055 (J.N.S., J.W.E.), W.W. Smith Charitable Trust Fund grant H1206 (J.N.S.) and University of Oxford, Radcliffe Department of Medicine pump-priming award (to J.N.S., Y.C., and P.M.H.). P.M.H. was additionally funded by a Royal Commission 1851 Research Fellowship, and Y.C. funded Alzheimer’s Research UK (ARUK-RF2019B-004). J.N.S and Y.C. were also supported by the Oxford BHF Centre of Research Excellence (RE/18/3/34214).

## Author contributions

J.N.S. conceptualized the study, with input from Y.C., P.M.H. and J.W.E. S.A. performed imaging experiments and function assays in mouse neuronal cells, N.J. performed imaging studies and TPC2 knockout validation in cardiomyocytes, K.M.C. performed calcium retention capacity assays, K.H., P.M.H. designed and performed experiments in human iPSC-induced neurons. Y.C., J.N.S. designed and performed experiments using the mouse tMCAO model. J.N.S, S.A, Y.C. wrote the manuscript with help and input from all authors. All authors contributed to data analysis and have read and approved the manuscript.

## Materials and Correspondence

Correspondence and requests for materials should be addressed to J.N.S.

## Abbreviations

Ca^2+^: calcium
OxPhos: oxidative phosphorylation
I/R: ischemia/reperfusion
ER: endoplasmic reticulum
TRPML1: transient receptor potential mucolipin 1
TPC2: two-pore channel 2
Na^+^: sodium
PI(3,5)P₂: phosphatidylinositol 3,5-bisphosphate
NAADP: nicotinic acid adenine dinucleotide phosphate
IP_3_R: inositol 1,4,5-trisphosphate receptor
mtCU: mitochondrial calcium uniporter channel
ΔΨm: mitochondrial membrane potential

## Extended data

**Extended Data Figure 1:**
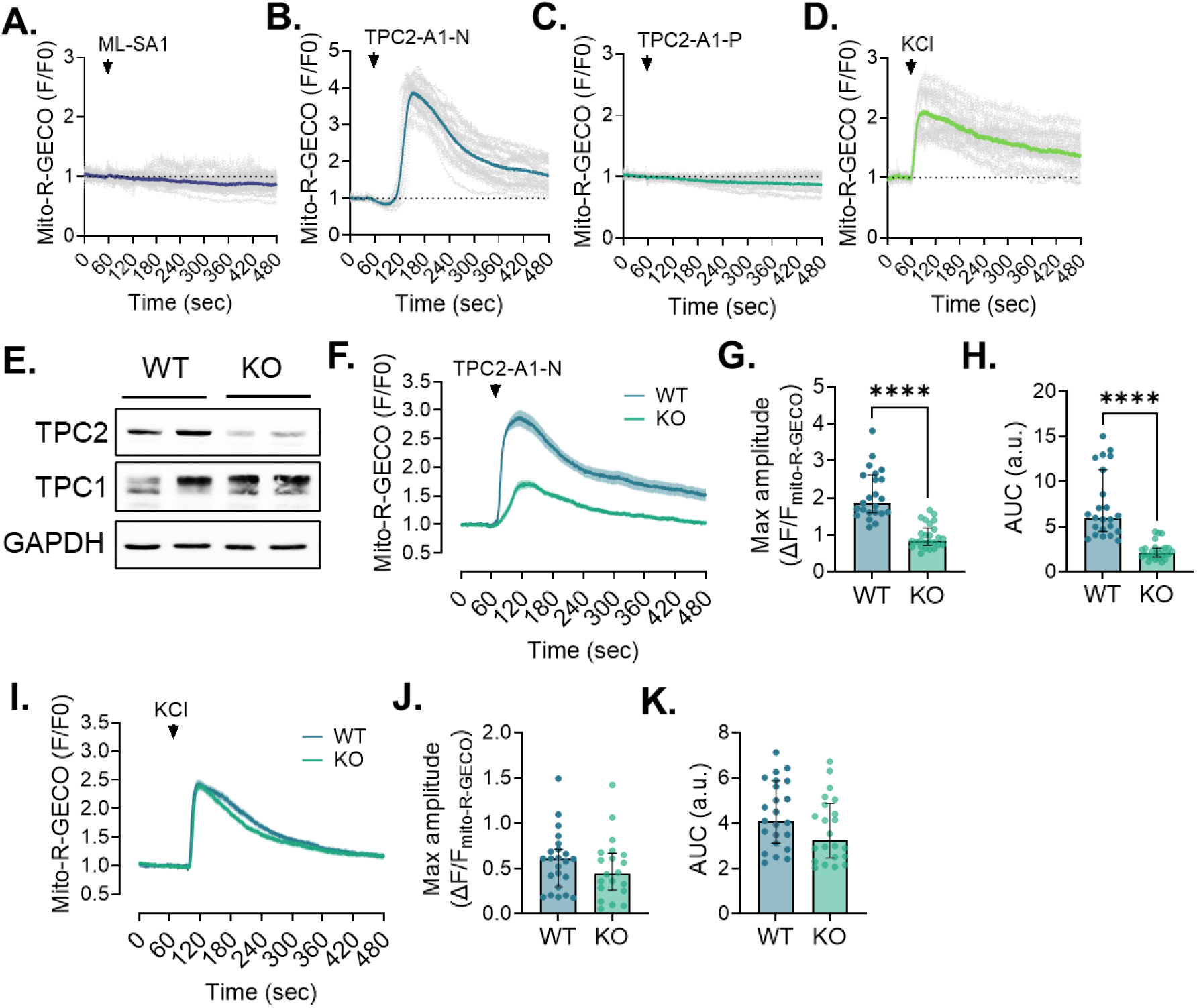
Cardiomyocytes display similar TPC2-dependent lysosome-mitochondrial Ca^2+^ communication. Averaged traces showing the change in mito-R-GECO intensity (normalized to baseline) in response to treatment with (**A**) ML-SA1 (30 μM, n=29), (**B**) TPC2-A1-N (30 μM, n=20), (**C**) TPC2-A1-P (60 μM, n=20), or (**D**) KCl (100 mM, n=14). Agonist-dependent responses were obtained on multiple cells across 3 separate days. (**E**) Immunoblotting of clonal AC16 cardiomyocyte lines after CRISPR–Cas9-mediated disruption of TPC2. (**F**) Averaged traces showing a significantly reduced mito-R-GECO response to TPC2-A1-N (30 μM) treatment in TPC2 KO cells compared to WT controls. Quantification of mito-R-GECO (**G**) peak amplitude and (**H**) area under the curve in WT and TPC2 KO AC16 cells after TPC2-A1-N treatment. Student’s t-test; ****p< 0.0001. n=23-26 cells/N=3 days per group. (**I**) Averaged traces and quantification of mito-R-GECO (**J**) peak amplitude and (**K**) area under the curve following KCl treatment demonstrate that the genotype-dependent differences in TPC2-A1-N response between WT and TPC2 KO cells was not due to compensatory changes in global Ca^2+^ handling. Student’s t-test. n=20-23 cells/N=3 days per group.

**Extended Data Figure 2:**
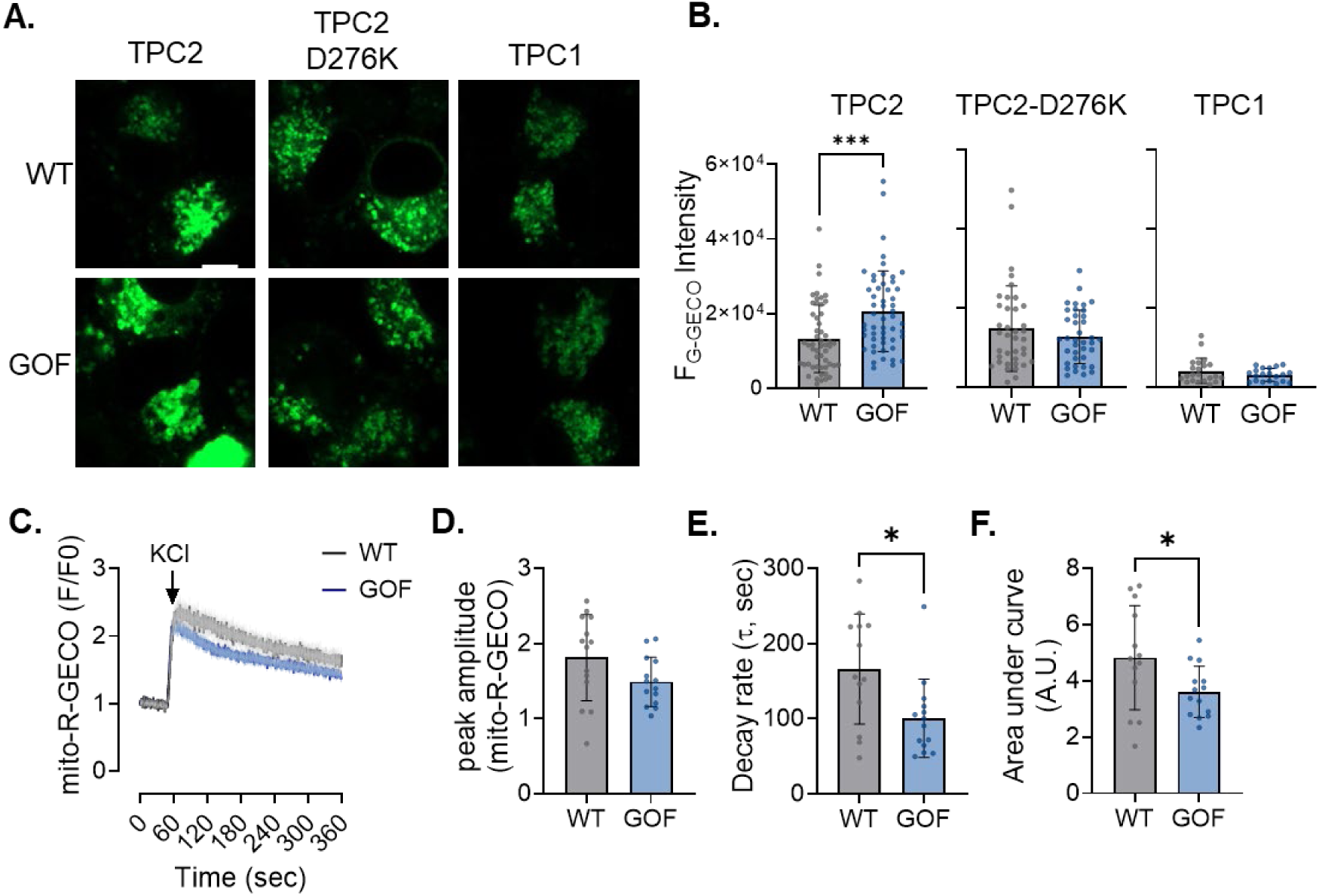
Validation of TPC2 GOF model. (**A**) Representative images of WT and TPC2 GOF N2A cells expressing either TPC2-G-GECO1.2 (wild-type channel), the pore-dead TPC2(D276K)-G-GECO1.2 mutant, or TPC1-G-GECO1.2. (**B**) Quantification of basal lysosomal Ca^2+^ release measured by G-GECO1.2, showing a modest but significant elevation in baseline TPC2-dependent Ca^2+^ efflux in TPC2 GOF cells compared with WT, whereas TPC2(D276K) and TPC1 constructs do not exhibit increased activity. Student’s t-test; ***p < 0.001. n = 23–25 cells per group. Scale bar = 5 µm. (**C**) Averaged traces of KCl–evoked mitochondrial Ca^2+^ transients in WT and TPC2 GOF N2A cells (n = 13 WT; n = 14 GOF). (**D–F**) Quantification of mitochondrial Ca^2+^ dynamics from (A), including peak amplitude (**D**), decay rate (**E**), and area under the curve (**F**). Student’s t-test; *p ≤ 0.05. N=3 independent days per group.

**Extended Data Figure 3:**
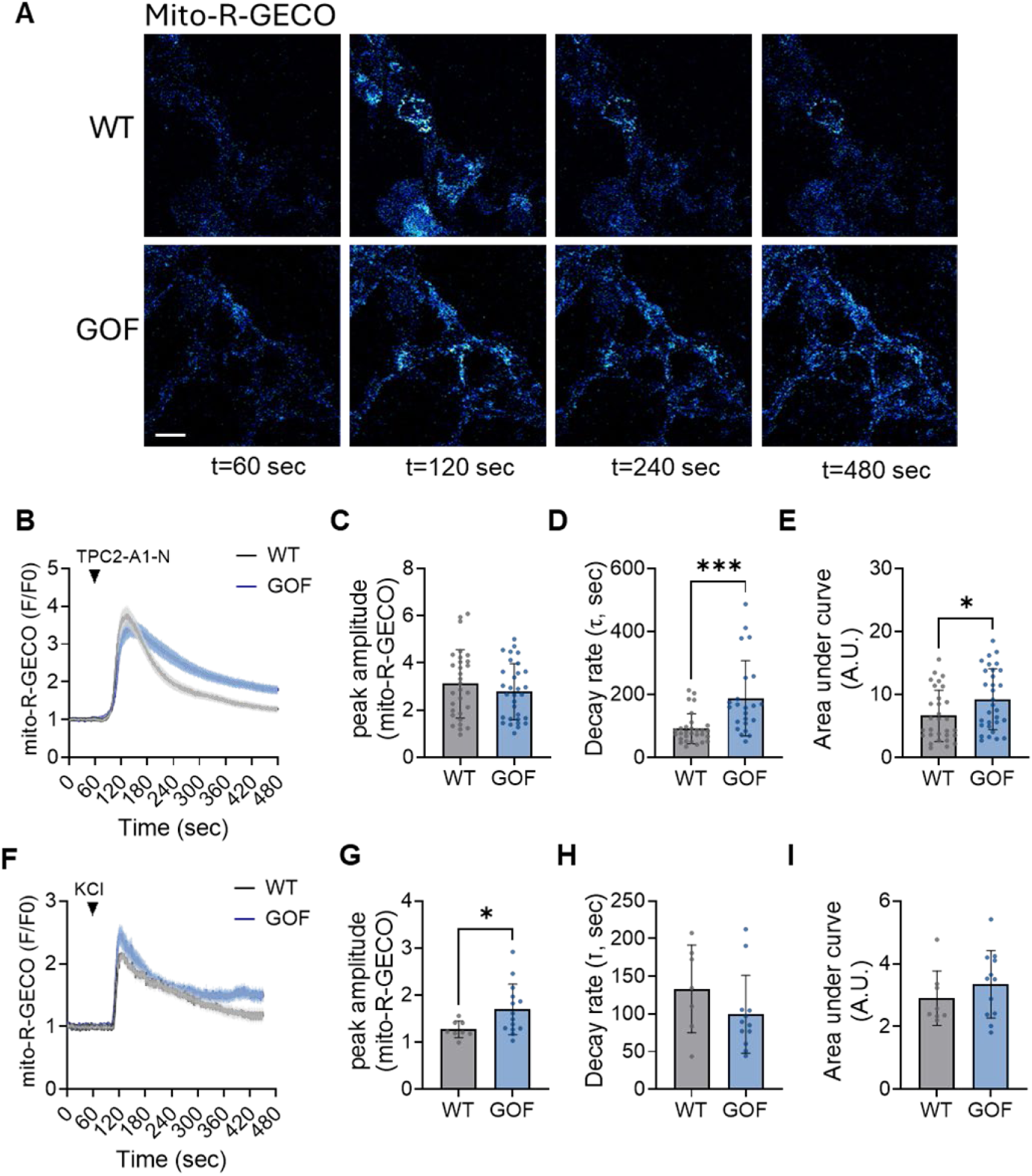
Cardiomyocytes show similar TPC2-dependent enhancement of mitochondrial Ca^2+^ load. (**A**) Representative time-lapse images of TPC2-A1-N–evoked mitochondrial Ca^2+^ signals (mito-R-GECO) in WT and TPC2 GOF AC16 cardiomyocytes (scale bars = 20 µm). (**B**) Averaged traces of TPC2-A1-N–evoked mitochondrial Ca^2+^ transients in WT and TPC2 GOF N2A cells (n = 30 WT; n = 31 GOF). (**C–E**) Quantification of mitochondrial Ca^2+^ dynamics from (A), including peak amplitude (**C**), decay rate (**D**), and area under the curve (**E**). (**F**) Averaged traces of KCl–evoked mitochondrial Ca²⁺ transients in WT and TPC2 GOF AC16 cardiomyocytes (n = 8 WT; n = 14 GOF). (**G–I**) Quantification of mitochondrial Ca^2+^ dynamics from (F), including peak amplitude (**G**), decay rate (**H**), and area under the curve (**I**). Student’s t-test; *p ≤ 0.05, ***p ≤ 0.001. N=3 independent days per group.

**Extended Data Figure 4:**
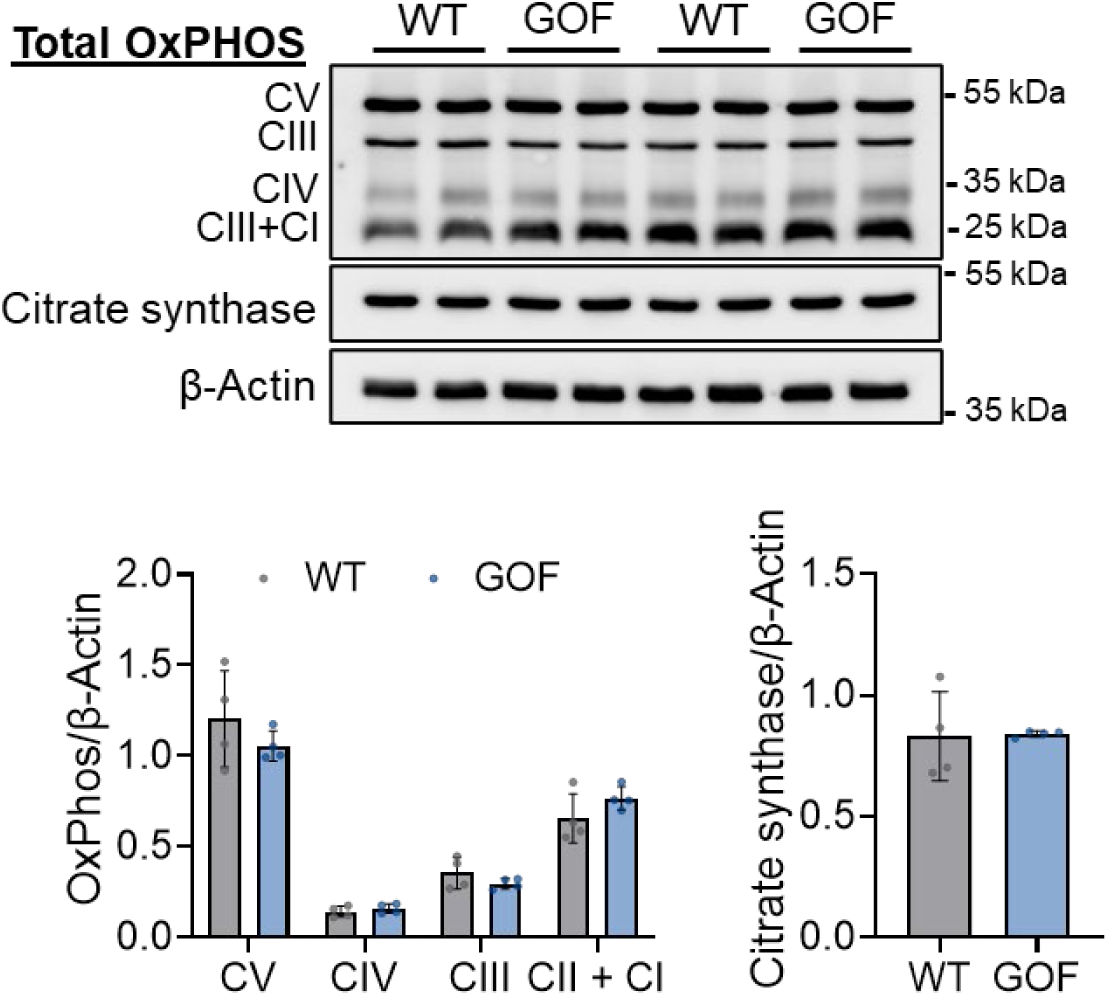
Comparable mitochondrial content between WT and GOF N2A cells. Representative western blot images and quantification of equal levels of ETC complex I-V, as evaluated using the pan-OxPHOS antibody and citrate synthase expression between WT and GOF N2A cells. Data normalized to the housekeeping protein, β-actin; n=4/grp.

